# High threat intensity increases conditioned fear generalization and makes extinction of a generalization stimulus less effective

**DOI:** 10.1101/2024.11.29.625989

**Authors:** Alba López-Moraga, Zeynep Gültekin, Laura Luyten, Tom Beckers

## Abstract

Generalization of conditioned fear is adaptive for survival. However, overgeneralization of fear from threat cues to loosely similar yet safe stimuli is a hallmark of anxiety-related disorders. Such overgeneralization may impact other fear learning processes. In particular, broad fear generalization might limit the effectiveness of extinction training, which may further maintain anxiety.

Here, we examined if increased generalization of conditioned fear might reduce the later generalization of fear extinction. To this end, we compared rats conditioned with moderate-versus high-intensity footshocks, the latter showing stronger generalization of acquired fear than the former. Within each shock intensity group, we then conducted extinction training using the original conditioned stimulus (CS) for half of the animals and a generalization stimulus (GS) for the others. We found stronger preserved responding to the CS after GS extinction in rats that had initially been conditioned using the high-intensity footshock than in rats conditioned using the moderate-intensity footshock, in line with the notion of reduced generalization of extinction from the GS to the CS in the former group. However, we also found indications for weaker retention of GS extinction in rats conditioned with the high-intensity footshock, which may in part or in whole explain the apparent difference in generalization of GS extinction to the CS.

Our results may be important to consider in extinction-based exposure therapy, where patients often present with broadly generalized fears and exposure treatment is usually not conducted using the exact cues or situations for which fear was initially acquired.

**Highlights:**

- We see greater CS-to-GS fear generalization using a strong (vs moderate) US in rats
- Paradoxically, we see more preserved CS fear upon GS extinction in the former group
- This may reflect an effect on extinction generalization and/or extinction retention
- CS extinction reduced GS fear similarly in both groups
- Male rats showed higher freezing during acquisition and extinction than females

## Introduction

Generalization of acquired fear is an adaptive phenomenon. In daily life, animals and humans may encounter threat-predictive stimuli in various forms and under various conditions, and learning to generalize from one cue for impending danger to another one can be vital for survival. However, overgeneralization, or the tendency to exhibit fear when confronted with essentially safe cues that merely resemble a genuine cue for danger, can interfere with adaptive functioning, and has been suggested to be a hallmark of anxiety and stress-related disorders (Cooper et al., 2022).

Generalization can be investigated experimentally through fear conditioning studies in humans, but thanks to a high degree of conservation across species in the laws and mechanisms that govern fear learning (Haaker et al., 2019), the fear conditioning paradigm can readily be applied in rodents as well, allowing to study generalization in a highly controlled environment and to investigate sex differences in the absence of sociocultural factors. Studies on fear generalization in rodents typically start with a fear acquisition phase, where a neutral stimulus (CS, e.g., a tone) is paired with an aversive footshock (US). Later, animals are presented with different cues that resemble the original CS (e.g., other tones). The degree of defensive responding that generalizes to those cues depends on how perceptually similar they are to the CS (Asok et al., 2019; Luyten et al., 2016; Zhang et al., 2017).

Prior research has shown that while fear acquired to a CS will generalize readily to a sufficiently similar generalization stimulus (GS), extinction training of such a GS, in which the GS is repeatedly presented without the US, will leave fear responding to the original CS or other GSs relatively intact in humans (Vervliet et al., 2004, 2005; but see Struyf et al., 2018) and rodents (Alfei et al., 2020, 2021; Boddez et al., 2012; Bouton and Bolles, 1979), whereas extinction training of the original CS will reduce fear responding to the CS as well as GSs.

Individuals with high trait anxiety show such lack of generalization of GS extinction to the original CS even more markedly (Wong and Lovibond, 2020). This observation has clinical relevance, because exposure therapy is often conducted using generalization cues (e.g., cars and traffic situations in general) rather than with the original cues that a patient acquired fear for (e.g., the car they had a crash in, the intersection they got run over at). In fact, as fear generalizes more broadly, chances increase that a therapist will expose a patient to generalized fear-eliciting situations – if they have become afraid of all sorts of traffic, it is all the more likely that a therapist will just expose them to a traffic situation that elicits strong fear, regardless of the exact nature of the initial incident.

It remains untested whether individuals who exhibit broader generalization of acquisition would show more of a reduction of defensive responding to the CS following extinction training with a GS. Formal models of associative learning suggest that the amount of generalization depends on the degree of perceived similarity between the CS and GS. Thus, strong generalization from the CS to a GS would imply that GS extinction should also generalize well to the CS (Pearce, 1987). As such, individuals who overgeneralize threat following fear acquisition should also show increased generalization of safety following extinction learning. However, a better-safe-than-sorry strategy would suggest that strong generalization of acquisition might in fact go hand-in-hand with *reduced* generalization of extinction, because incorrectly identifying a dangerous stimulus as safe is more costly than incorrectly identifying a safe stimulus as dangerous (Dunsmoor and Paz, 2015). This asymmetry might prevent especially cautious individuals, who would show stronger generalization from the CS to a GS, from generalizing extinction learning for that GS to the original fear-eliciting CS or to other GSs (Beckers et al., 2023). Here, we studied whether heightened generalization of fear after acquisition would indeed weaken the later generalization of fear extinction.

In the present study, we performed fear conditioning using moderate- and high-intensity footshock USs in male and female rats, in light of the evidence that high-intensity USs promote stronger fear generalization than moderate-intensity USs (Laxmi et al., 2003; Xuan et al., 2023). Subsequently, we conducted fear extinction using either the CS or a GS. We hypothesized that enhanced generalization of fear from CS to GS after acquisition would weaken the later generalization of fear extinction from GS to CS. Additionally, we expected female rats to show overall slower extinction than male rats (Baran et al., 2009; Fenton et al., 2016; Greiner et al., 2019) and more generalization of freezing from CS to GS (Greiner et al., 2019; Keiser et al., 2017).

## Methods

### Preregistration and data availability

The design, procedures, sample size and analysis plan were preregistered on the Open Science Framework (OSF) prior to the start of data collection (http://doi.org/10.17605/osf.io/9yd83). All data and scripts are available through the same OSF link as well.

### Subjects

The experiment was performed in accordance with Belgian and European laws (Belgian Royal Decree of 29/05/2013 and European Directive 2010/63/EU) and the ARRIVE 2.0 guidelines (Percie du Sert et al., 2020) and approved by the KU Leuven animal ethics committee (project license number 011/2019). The study was conducted in 64 8-week-old male (270-300 g at arrival) and female (200-230 g at arrival) rats (Janvier Labs, France). Animals were housed in groups of 4, on a 12-hour light-dark cycle (lights on at 7 am); experimental procedures were performed between 9 am and 5 pm. The cages had bedding and cage enrichment in the form of a hanging red polycarbonate tunnel. Water and food were available ad libitum for the entire experiment, except during behavioral testing. Animals were acclimated for one week to our facility and habituated to handling for 2 days before the start of the experiment.

### Apparatus

Eight identical operant chambers (30.5 cm width, 25.4 cm depth and 30.5 cm height; Rat Test Cage, Coulbourn Instruments, Pennsylvania, USA) were used simultaneously and were enclosed in sound-attenuating boxes. The shock grid consisted of 18 stainless steal rods of 0.4 cm diameter, with 1.5 cm spacing between the center of the rods (Modular Shock Floor H10-11R-TC-SF, Coulbourn Instruments). This grid was connected to a scrambled shocker (Precision Animal Shocker H13-15, Coulbourn Instruments). The different tones were generated by a modified tone generator (Seven-tone generator H12-07, Coulbourn Instruments) and transmitted to the operant chamber through two speakers in opposite walls of the chamber. The experiment was conducted using Graphic State 4 (Coulbourn Instruments) software with the house light off. Behavior was continuously recorded during the experimental sessions using an IP camera (Foscam C2M, Shenzhen, China) attached to the ceiling of the sound-attenuating box. The operant chambers were cleaned between animals with soapy water (5.55% Instanet, Henkel, Vilvoorde, Belgium).

### Experimental design

Rats (n = 64) were evenly divided into four groups: moderate shock during fear conditioning, extinction training with CS (MSCS), moderate shock during fear conditioning, extinction training with GS (MSGS), strong shock during fear conditioning, extinction training with CS (SSCS), strong shock during fear conditioning, extinction training with GS (SSGS) (see Figure 1). Rats of both sexes were divided equally over the four groups, the final group sizes were: MSCS (n = 16; 8 females), MSGS (n = 16; 8 females), SSCS (n = 16; 8 females) and SSGS (n = 16; 8 females). All sessions were scheduled 24h apart, except for the spontaneous recovery test, which was conducted 7 days after the reinstatement test.

**Figure 1.**
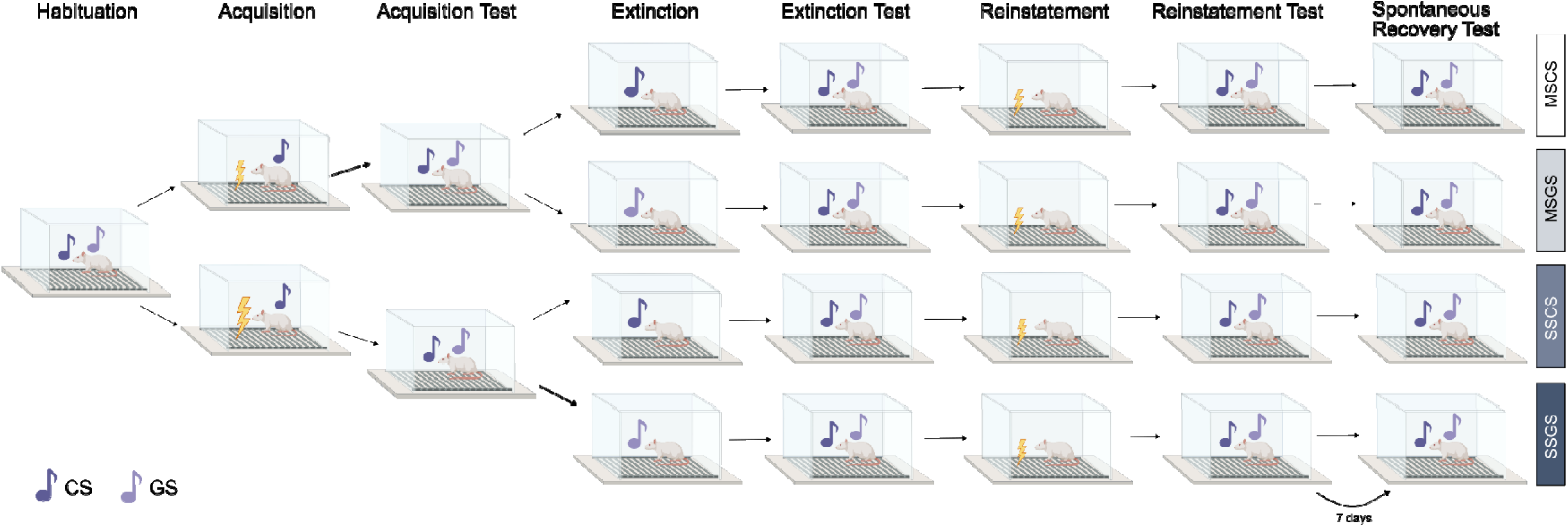
Experimental design. MSCS: moderate shock acquisition, extinction training with CS (n = 16); MSGS: moderate shock acquisition, inction training with GS (n = 16); SSCS: strong shock acquisition, extinction training with CS (n = 16); SSGS: strong shock acquisition, inction training with GS (n = 16). All sessions were scheduled 24h apart, except for the spontaneous recovery test, which was conducted 7 ys after the reinstatement test. At the end of extinction test, some rats were euthanized for tissue collection. Thus, in reinstatement and nstatement test all groups had n = 12, except the SSCS group which had n = 13 (n = 7 females, n = 6 males). After reinstatement test, some s were euthanized for tissue collection and sample sizes for spontaneous recovery test were n = 10, except the SSCS group which had n = (females n = 6, males n = 5) and the MSCS group which had n = 9 (n = 4 females, n = 5 males).

First, all rats underwent a habituation session during which the CS and GS were each presented twice. Half of the rats received the tone presentations in the sequence CS, GS, CS, GS, while the other half received them in the sequence GS, CS, GS, CS. The CS and GS consisted of a pure 3-kHz tone and a pure 7-kHz tone, counterbalanced between animals. After 3 min of acclimation, each tone was presented for 30 s, with an average ITI of 300 s (270-330 s). After the last tone, the rats remained in the testing chamber for one additional minute.

The next day, rats underwent an acquisition session. After 3 min of acclimation, 9 CS-US pairings were presented, with an ITI averaging 180 s (150-210 s). Half of the rats (n = 32) were trained using a moderate footshock US of 0.4 mA intensity and 1 s duration. The other half (n = 32) were trained using a strong footshock US of 1 mA intensity and 2 s duration. The 30-s CS co-terminated with the US in both conditions. After the last CS-US pairing, rats remained in the testing chamber for an additional minute. One day later, an acquisition retention test was performed, during which the CS and GS were each presented twice, with the same parameters as in the habituation session.

The extinction session consisted of 18 CS or GS presentations, after 3 min of acclimation, and with an ITI averaging 180 s (150-210 s). After the last tone presentation, rats remained in the testing chamber for an additional minute. Four rats per group, 2 of each sex, were perfused 90 min after the extinction test to dissect their brains for immunohistochemical staining. The next day, an extinction retention test was performed, during which the CS and GS were each presented twice, with the same parameters as in the habituation session.

A reinstatement manipulation took place one day later. After 3 min of acclimation, a single footshock of 0.4 mA and 1 s duration was delivered, followed by 2 more minutes in the testing chamber. The next day, a reinstatement test session was conducted, consisting of 2 CSs and 2 GSs, as in the habituation session. Two rats per group, one of each sex, were perfused 90 min after the reinstatement test.

Seven days after the reinstatement test, a spontaneous recovery test took place with 2 CSs and 2 GSs, using the same parameters as in the habituation session. Four rats per group, two of each sex, were perfused 90 min after the spontaneous recovery test.

Due to problems with tissue quality after freezing the specimens, the planned immunohistochemical stainings were not performed.

### Behavior

Freezing during CS presentations was scored manually from the videos as a percentage of CS duration. Freezing was defined as full immobility except for minimal movements associated with breathing.

A freezing discrimination index was calculated for sessions where both CSs and GSs were presented. It was calculated as the difference in freezing between the two different tones (i.e., CS minus GS) divided by the sum of freezing to both tones (Xu and Südhof, 2013).

#### Automated analyses of behavior

To investigate darting behavior and contextual freezing, we used a combination of DeepLabCut (version 2.3.9) (Mathis et al., 2018; Nath et al., 2019) and SimBA (version 1.99.5) (Goodwin et al., 2024).

First, we used DeepLabCut to obtain pose estimation. We labeled 200 frames taken from 10 videos, each video coming from a different animal; 95% of these frames were used for training. Five body parts were selected: nose, left ear, right ear, centroid, and tail base. We used a RestNet-50 neural network with default parameters for 300000 training iterations. Given that our results were initially unsatisfactory (see OSF for further details), we extracted outliers, relabeled 20 of those additional outlier frames for each video and ran an additional training iteration with the same parameters as above. We validated with 1 shuffle and found that the test error was 1.8 pixels and the training error was 5.78 pixels (image size: 456 x 256 pixels, downsampled from original 1280 x 720 pixels).

Second, we used SimBA to further analyze pose estimation data obtained with DeepLabCut. We created a single animal project configuration with user-defined body parts (same body parts as for DeepLabCut). We defined pixels per mm by considering the distance between the width of the bottom of the fear conditioning chamber, which in our case was 291 mm. We used the body-part: nearest interpolation method and a gaussian smoothing method of 100 ms, outlier correction was skipped.

To analyze darting, we obtained velocity traces per animal at 1-s time bins. We filtered speeds under 23.5 cm/s. An animal was considered a darter when a darting bout occurred during CS 3 to 7 of the acquisition session, excluding intertrial intervals and US presentations (Gruene et al., 2015).

To analyze contextual freezing, we started from a model that we described and used previously (López-Moraga et al., 2025). We annotated three additional videos; we added 972 frames (32.4 s) annotated with freezing present and 24491 frames (816.33 s) annotated with freezing absent. Here, we specifically focused on labelling sniffing as a non-freezing bout. We built the model with the following settings: n_estimators = 2000, RF_criterion = gini, RF_max_features = sqrt, RF_min_sample_leaf = 1, under_sample_ratio = 3, class_weights = custom: Freezing present = 2; Freezing Absent = 1, with an 80% training split. The evaluation performance of the freezing classifier had an F1 score of 0.877, precision of 0.876, and recall of 0.878 for freezing present and F1 score of 0.960, precision of 0.960, and recall of 0.959 for freezing absent. We set the discrimination threshold at Pr = 0.75 and a minimum duration of 500 ms (see OSF for further details).

### Data analyses

Statistical analyses were preregistered; any deviations from the preregistered analyses are specified below. Data were analyzed using t-tests and one-way, two-way, repeated-measures or mixed ANOVAs, as appropriate. Post-hoc tests with corrections for multiple testing followed if significant main effects or interactions were found. If assumptions were violated, nonparametric statistical analyses were performed. For the nonparametric mixed RM ANOVAs, we applied an aligned rank transform ANOVA using the ARTool package (Elkin et al., 2021) in R (R Core Team, 2022) and R Studio (RStudio Team, 2021). All other statistical analyses were performed using performance (Lüdecke et al., 2021), afex (Singmann et al., 2023), emmeans (Lenth et al., 2024) and effectsize (Ben-Shachar et al., 2024) R packages. Data were processed and plotted using the tidyverse (Wickham and RStudio, 2022) R packages.

Analyses for habituation and acquisition sessions were not preregistered. Habituation was analyzed with a non-parametric mixed ANOVA with shock intensity (moderate, strong) as a between-subjects variable and stimulus (CS, GS) as a within-subjects variable. Acquisition performance was analyzed with a two-way ANOVA with shock intensity and sex as between-subjects factors.

Data from the acquisition retention test, extinction retention test, reinstatement test and spontaneous recovery test were analyzed with sex and either shock intensity (moderate, strong) or experimental group (MSCS, MSGS, SSCS, SSGS) as between-subjects factors and stimulus (CS, GS) as within-subjects factor. If effects of sex were not significant, both sexes were collapsed in the mixed ANOVA as preregistered. As a deviation from the preregistration, we did not directly compare retention test sessions (i.e., acquisition test versus extinction test) within one mixed ANOVA. Two-tailed one-sample t-tests were performed on the discrimination indices for the acquisition test and extinction test of each shock intensity group, comparing the mean of each group to 0 (e.g., no discrimination). This comparison was not preregistered. The analysis of contextual freezing during the extinction retention test was exploratory and consisted of a mixed ANOVA with sex and group as between-subject variables and minute as within-subjects variable.

Differences in freezing during extinction training were analyzed using a mixed ANOVA with trial as a within-subjects factor and sex and experimental group as between-subjects factors. In a separate mixed ANOVA, to assess changes in freezing, the first three tones (i.e., early block) were compared to the last three tones (i.e., late block) using block as a within-subjects factor and group and sex as between-subjects factors.

Criteria for classifying a rat as darter or non-darter were preregistered as described above. The chi-square test to compare the ratio of darters between sexes was preregistered, but the analyses comparing the ratio of darters between groups were exploratory. The analysis of differences in freezing between darters and non-darters by experimental session and sex were preregistered, except for the analysis of the extinction test session. The correlation analysis between number of darting bouts and freezing was exploratory.

## Results

The aim of this experiment was to investigate whether increased generalization of acquisition from CS to GS would go hand in hand with reduced generalization of extinction from GS to CS. To evaluate this, we needed to manipulate the generalization of acquisition. We therefore divided our sample in two groups, one that received a moderate intensity footshock as US during acquisition and one that received a strong intensity footshock as US, as higher US intensity has been shown to promote increased conditioned fear generalization (Dunsmoor et al., 2017; Laxmi et al., 2003; Xuan et al., 2023). After a test of generalization of acquisition, we conducted extinction training with either the CS or a GS and performed an extinction retention test 24 h later to test our central hypothesis. Finally, we performed exploratory tests of reinstatement and spontaneous recovery.

### Manipulation checks

In the habituation session, all rats exhibited uniformly low levels of freezing (F(1, 62) = 1.58, p = 0.213, ω_p_^2^ < -0.01, see further analyses in Table 1), with no significant differences in discrimination between CS and GS (t(31) = 0.65, p = 0.520, d = 0.12) (see Figure 2).

**Table 1.**
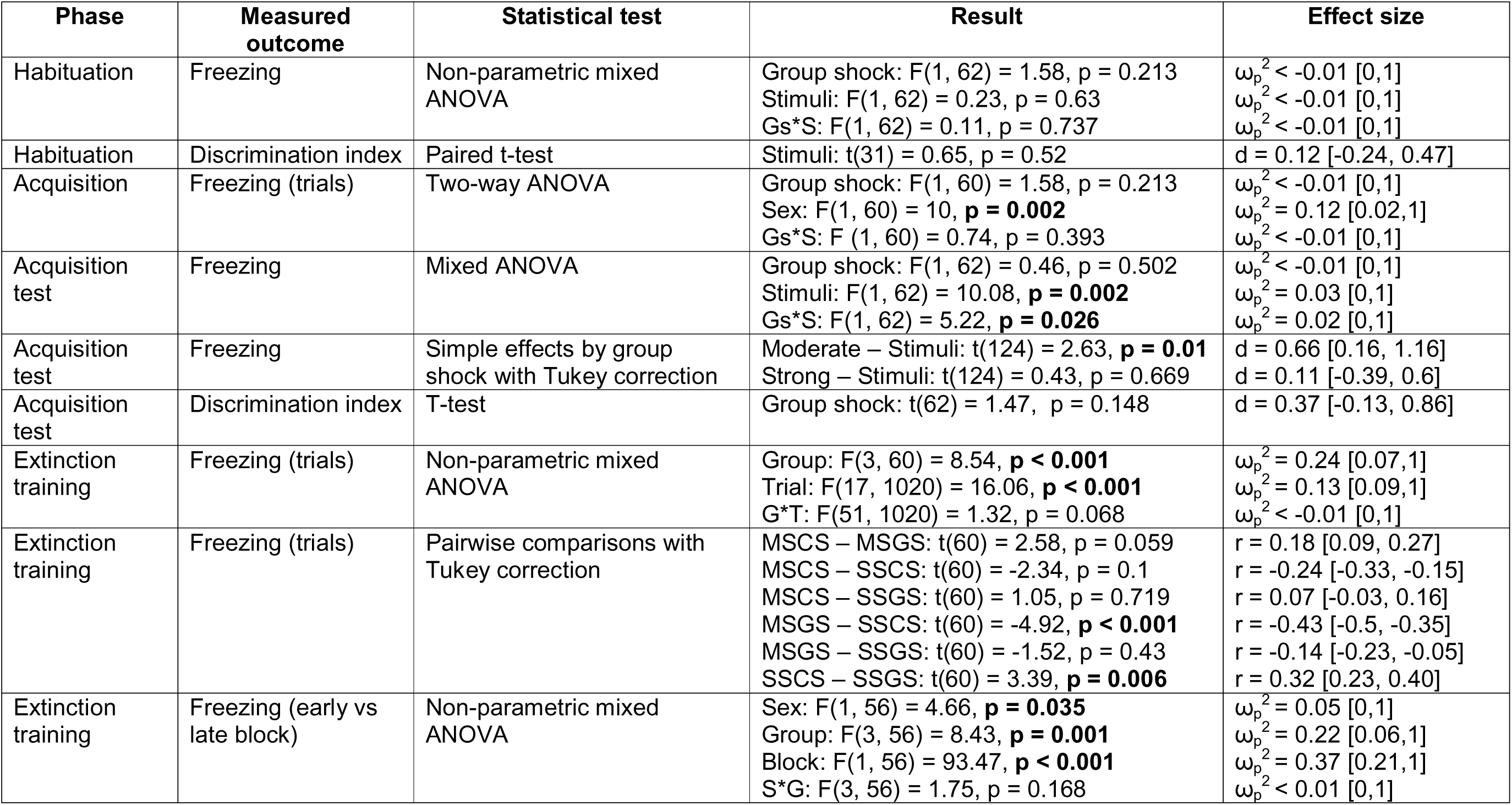
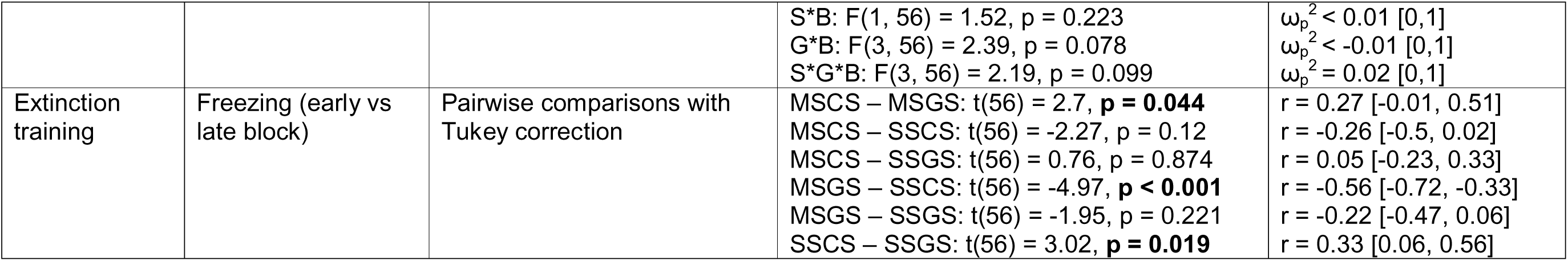
Statistical test results of the manipulation check.

**Figure 2.**
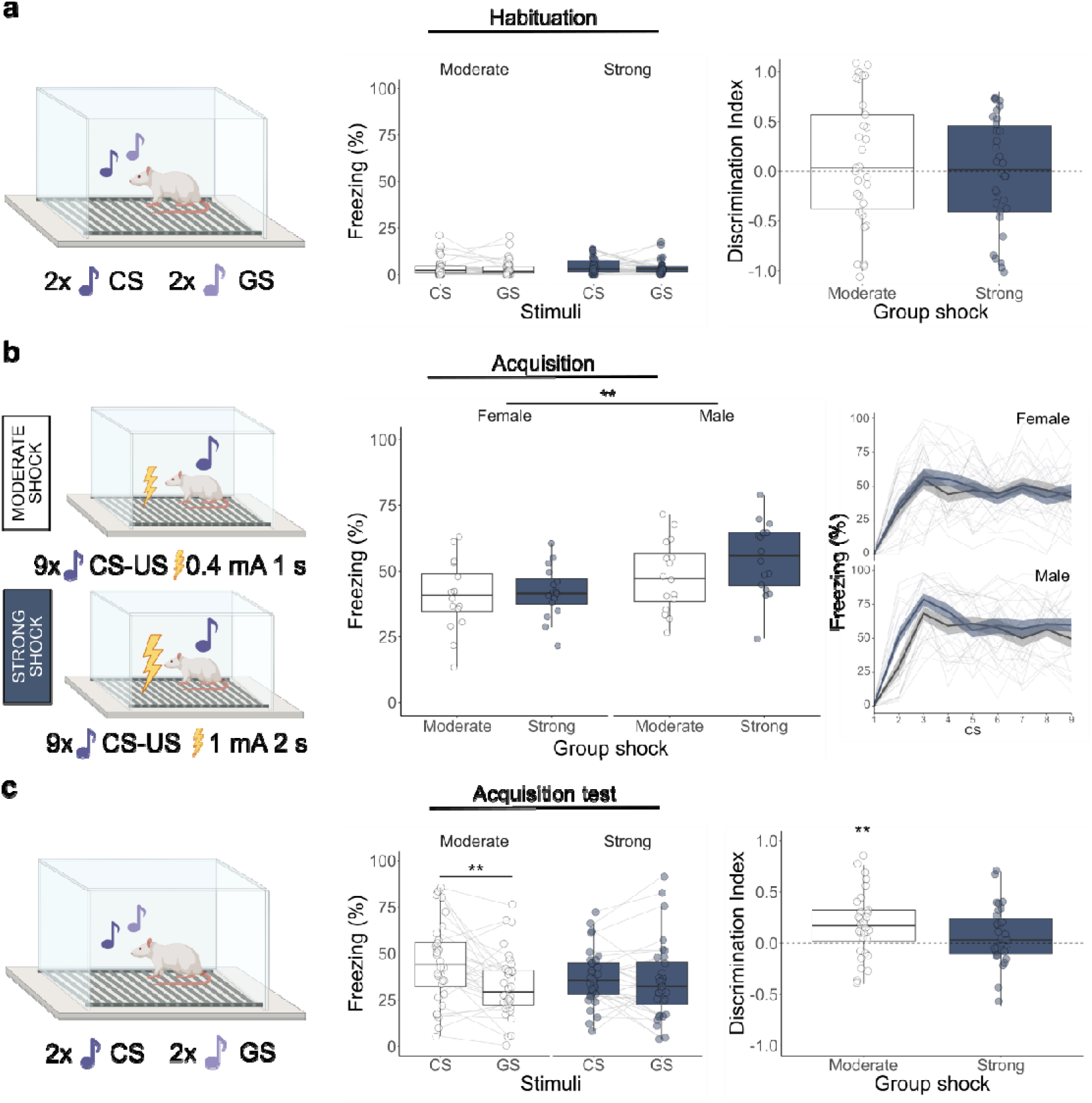
Freezing to the tones during habituation, acquisition, and the acquisition retention test and discrimination indices during habituation and acquisition test. Freezing boxplots represent average freezing to each stimulus, expressed as % of tone duration. Discrimination boxplots represent the average freezing discrimination index per group. Discrimination indices were calculated as the difference in freezing during the presentation of the CS versus the GS, divided by the sum of freezing to both tones. **a.** Graphical representation and results of the habituation session. **b.** Graphical representation and results of the acquisition session. Males showed higher freezing than females during acquisition (F(1, 60) = 10, p = 0.002). In the line plot bold lines represent the mean and the surrounding shaded area represents the standard error of the mean. **c.** Graphical representation and results of the acquisition retention test. An interaction between shock intensity group and freezing (F(1, 62) = 5.22, p = 0.026) reflects the fact that there were significant differences in freezing between CS and GS in the moderate shock group (t(124) = 2.63, p = 0.01), but not in the strong shock group (t(124) = 0.43, p = 0.669). The discrimination index was significantly greater than 0 for rats in the moderate shock group (t(31) = 3.34, p = 0.002), but not the strong shock group (t(31) = 1.32, p = 0.196).

**Table 2.**
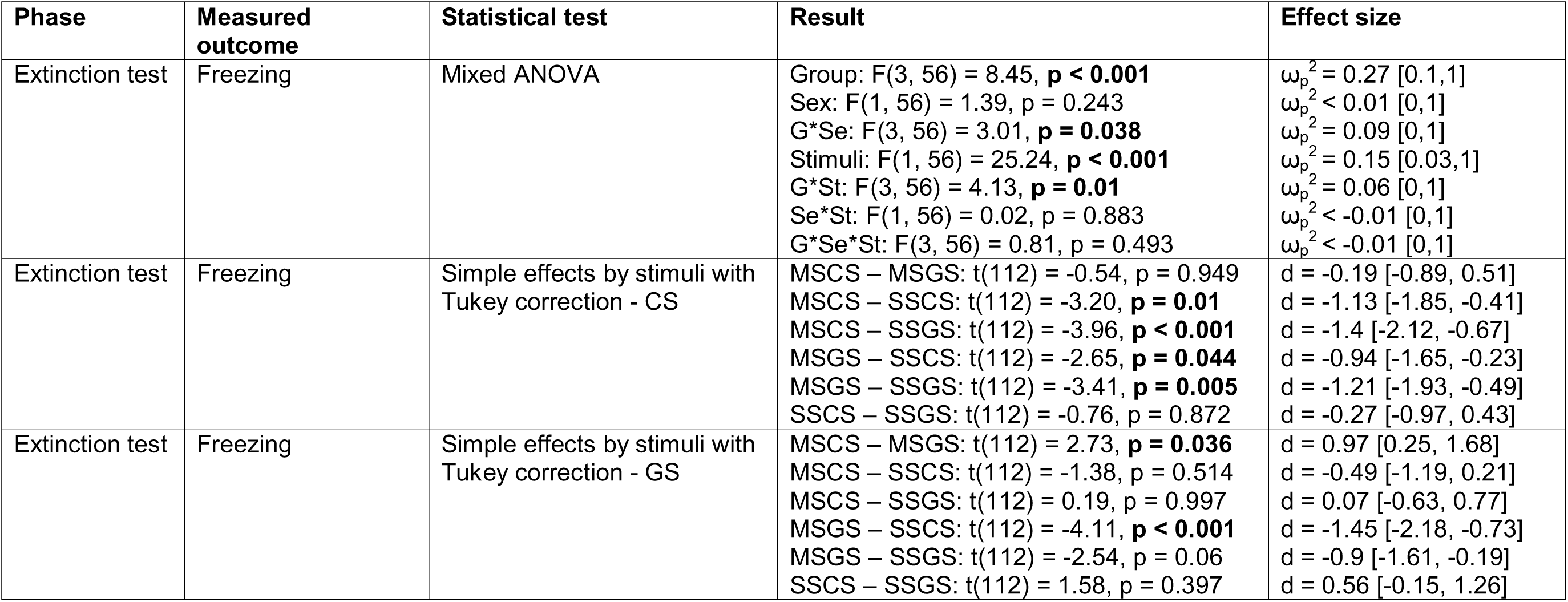
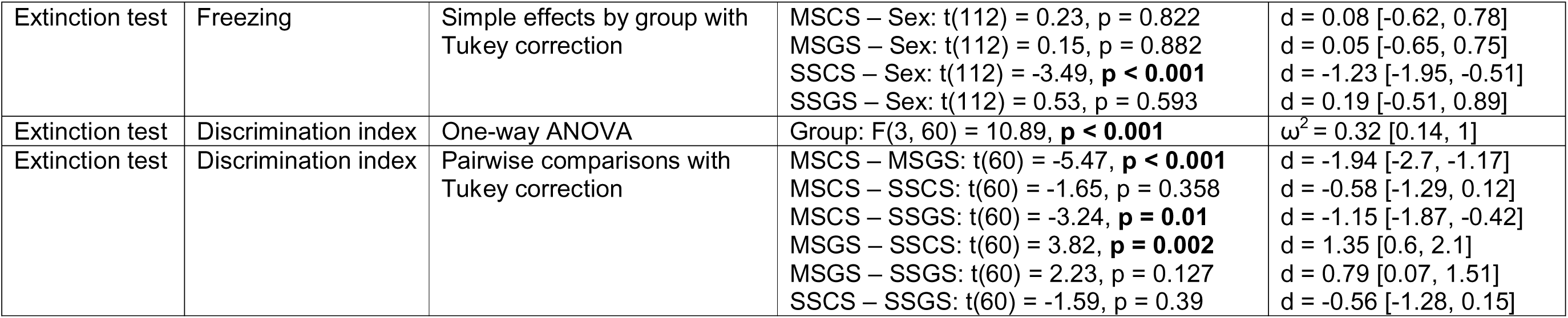
Statistical test results for the effects of generalized acquisition on generalization of extinction.

**Table 3.**
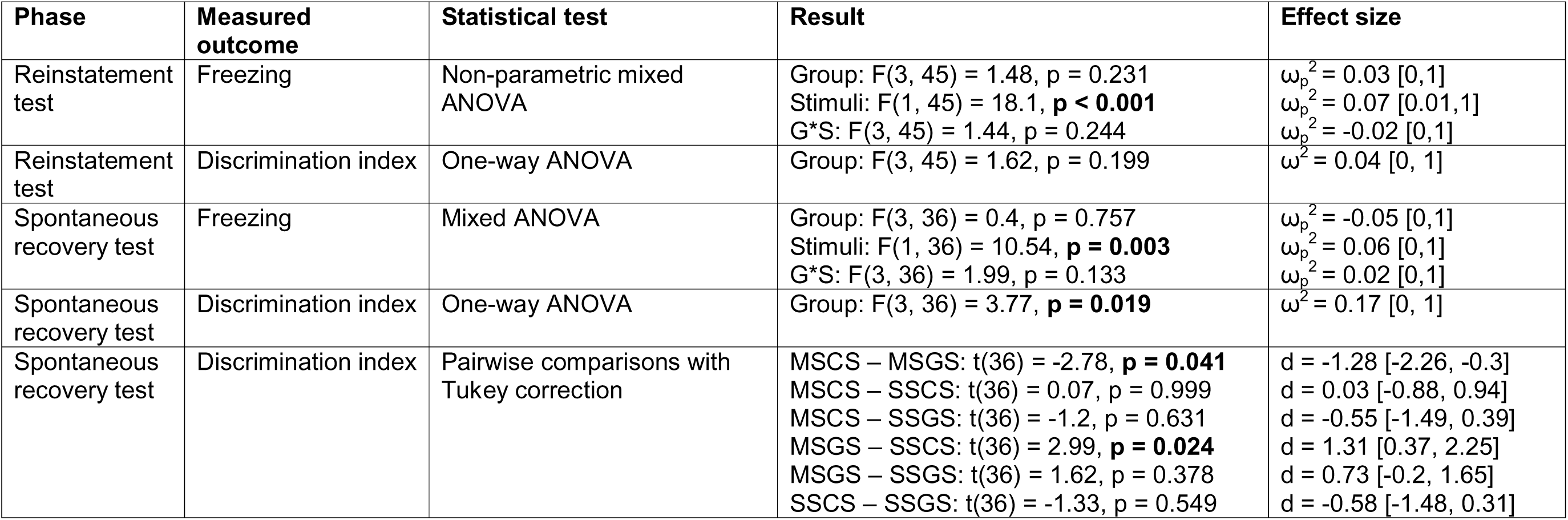
Statistical tests for return of fear.

During acquisition, we did not observe differences in freezing between rats that experienced moderate footshock USs or strong footshock USs (F(1, 60) = 1.58, p = 0.213, ω_p_^2^ < -0.01), but males showed significantly higher freezing than females (F(1, 60) = 10.00, p = 0.002, ω_p_^2^ = 0.12).

We also investigated darting behavior during acquisition. Rats were classified as darters or non-darters according to previous literature (Gruene et al., 2015). Forty-seven percent of female rats (15/32 rats) were classified as darters, against 19% (6/32) of male rats. A chi-square test revealed a significant association between darter status and sex (χ^2^(1)= 4.54, p = 0.033, φ = 0.3, see Figure S1A). This suggests that the distribution of darters was different between sexes. Additionally, female darters performed more dart bouts (M = 3.47, SE = 0.68) than male darters (M = 1.83, SE = 0.31) during acquisition. We did not find a significant association between shock intensity group and darter status (χ^2^(1)= 0.28, p = 0.594, φ = 0.1).

Given the different distribution of darters between sexes, we analyzed by sex whether there were differences in freezing during acquisition between darter and non-darter rats (Figure S1B). Female darters showed significantly less freezing than female non-darters (t(30) = - 4.8, p < 0.001, d = -1.7). However, in male rats, darting did not make a difference (t(30) = - 0.34, p = 0.73, d = -0.17). We did not find differences in freezing between darters and non-darters in later sessions, regardless of sex (see Supplementary Table 1 for further statistics).

In the acquisition test, there was a significant interaction between shock intensity group and stimulus (F(1, 62) = 5.22, p = 0.026, ω_p_^2^ = 0.02, see further analyses in Table 1). Specifically, the moderate shock group showed significantly higher freezing to the CS than the GS (t(124) = 2.63, p = 0.01, d = 0.66), whereas the strong shock group showed no significant difference in freezing between the stimuli (t(124) = 0.43, p = 0.669, d = 0.11), suggesting stronger generalization in the latter group, although the freezing discrimination index did not differ significantly between both groups (t(62) = 1.47, p = 0.148, d = 0.37). Additionally, we performed an exploratory one-sample t-test to assess if the discrimination index was significantly different from 0 in each group. For rats in the moderate shock group, the discrimination index was significantly greater than 0 (t(31) = 3.34, p = 0.002, d = 0.59), whereas for rats in the strong shock group it was not significantly different from 0 (t(31) = 1.32, p = 0.196, d = 0.32).

For the subsequent extinction training phase, both shock intensity groups were to be split in two, with half of the rats receiving extinction training for the CS and the other half receiving extinction training for the GS. We checked whether any differences between the four groups-to-be were present already in the acquisition test (see Figure S2 and Table S2).

In extinction training, we observed a reduction of freezing over trials (F(17, 1020) = 16.06, p < 0.001, ω_p_^2^ = 0.13), and overall differences in freezing levels between groups (F(3, 60) = 8.54, p < 0.001, ω_p_^2^ = 0.24) (see Figure 3). In particular, the SSCS group exhibited significantly higher levels of freezing than the SSGS (t(60) = 3.39, p = 0.006, r = 0.32) and MSGS groups (t(60) = 4.92, p < 0.001, r = 0.43).

**Figure 3.**
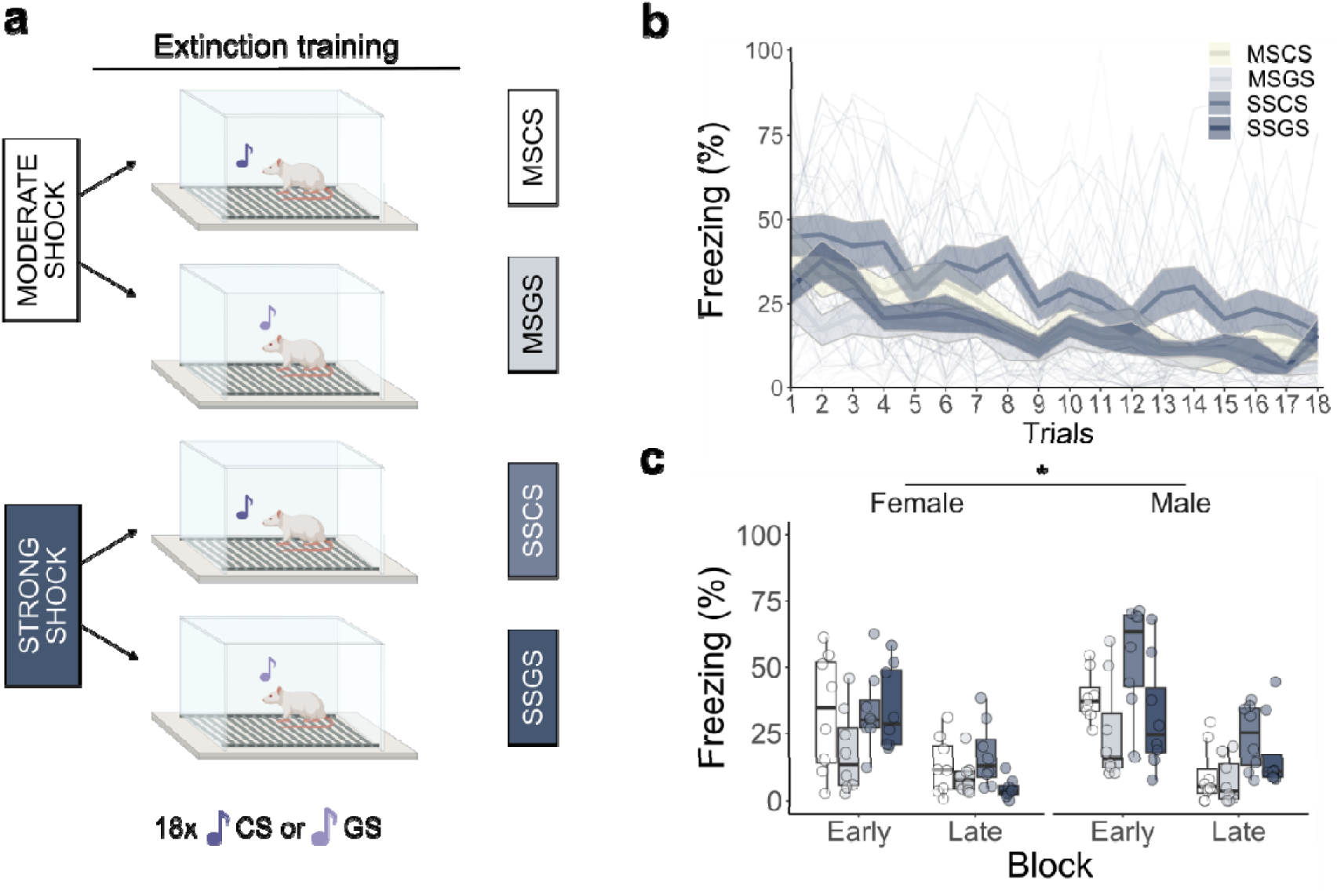
Freezing during extinction training, expressed in % of tone duration. **a.** Graphical representation of the extinction training session. **b.** Bold lines represent the mean and the surrounding shaded area the standard error of the mean. A reduction in freezing behavior was observed during extinction training (F(17, 1020) = 16.06, p < 0.001), along with significant differences between groups (F(3, 60) = 8.54, p < 0.001). **c.** Boxplots represent the average of the first three (i.e., early) and last three (i.e., late) tone presentations of the extinction session. While a general reduction in freezing was observed between the early and late block (F(1, 56) = 93.47, p < 0.001), males had higher freezing levels than females (F(1, 56) = 4.66, p = 0.035) and significant differences between groups were found (F(3, 56) = 8.43, p = 0.001).

Comparing freezing levels between an early block and a late block of the extinction training session, we found that males froze significantly more than females (F(1, 56) = 4.66, p = 0.035, ω_p_^2^ = 0.05) and that freezing was lower in the late block than in the early block, as expected (F(1, 56) = 93.47, p < 0.001, ω ^2^ = 0.37). Additionally, significant differences in freezing levels were detected between groups (F(3, 56) = 8.43, p = 0.001, ω_p_^2^ = 0.22). In particular, freezing levels differed between the two moderate shock groups, with rats undergoing extinction with the CS (MSCS) exhibiting higher freezing than rats undergoing extinction with the GS (MSGS) (t(56) = 2.7, p = 0.044, r = 0.27). Similarly, for the strong shock groups, SSCS rats showed higher levels of freezing than SSGS rats (t(56) = 3.02, p = 0.019, r = 0.33). We evaluated differences in freezing depending on sex and whether animals had been classified as darters during acquisition. We did not find significant differences between female darters and non-darters (t(30) = -1.37, p = 0.181, d = -0.49, see Figure S1C) nor between male darters and non-darters (t(30) = -1.3, p = 0.202, d = -0.64). 15

### Effects of generalization of acquisition on generalization of extinction

In the extinction retention test session, we observed a group by sex interaction (F(3, 56) = 3.01, p = 0.038, ω ^2^ = 0.09) and a group by stimulus interaction (F(3, 56) = 4.13, p = 0.01, ω_p_ = 0.06) (see Figure 4). Further investigation of the group by stimulus interaction, zooming in on the degree of generalization of extinction to the stimulus *not* presented during extinction training, revealed that rats in the SSGS group showed significantly higher freezing to the CS than rats in the MSGS group (t(112) = 3.41, p = 0.005, d = 1.21), whereas rats from the SSCS group and rats from the MSCS group did not display significant differences in freezing to the GS (t(112) = 1.38, p = 0.514, d = 0.49). Rats in the SSGS group had not shown higher freezing to the CS than rats in the MSGS groups in the acquisition test. Do note, however, that rats in the MSGS group also tended to show better extinction retention for the GS than rats in the SSGS group, even if not significantly so (t(112) = -2.54, p = 0.06, d = -0.9).

**Figure 4.**
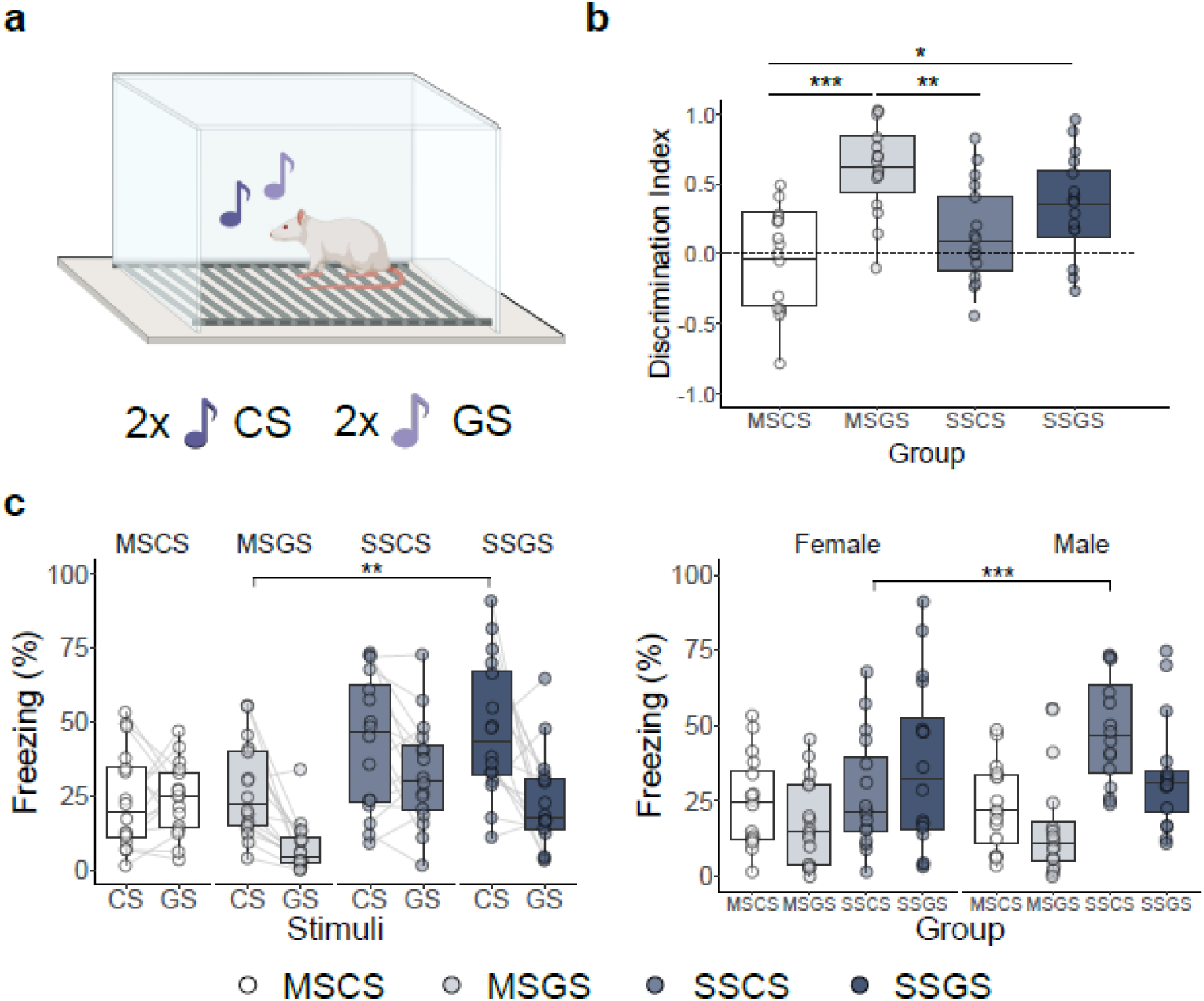
Results of the extinction retention test session. **a.** Graphical representation of the extinction retention test. **b.** The discrimination index boxplot represents the average of the freezing discrimination index. Discrimination indices were calculated as the difference in freezing to the different tones (CS versus GS), divided by the sum of freezing to both tones. Significant differences in discrimination were observed between groups (F(3, 60) = 10.89, p < 0.001). **c.** The freezing boxplots represent the average percentage of freezing to each stimulus, expressed in % of tone duration. Interactions between group and sex (F(3, 56) = 3.01, p = 0.038) and between group and stimulus (F(3, 56) = 4.13, p = 0.01) were found in freezing behavior. Rats in the MSGS group showed significantly lower freezing to the CS than rats in the SSGS group (t(112) = -3.41, p = 0.005). In the SSCS group, males showed higher freezing than females (t(112) = -3.49, p < 0.001).

Importantly, rats in the MSCS group did show significantly less freezing to the CS than those in the SSCS group (t(112) = -3.20, p = 0.01, d = -1.13). Group differences were detected in the freezing discrimination index as well (F(3, 60) = 10.89, p < 0.001, ω2 = 0.32). The MSCS group showed a significantly lower discrimination index than the MSGS group (t(60) = -5.47, p < 0.001, d = -1.94) and the SSGS group (t(60) = -3.24, p = 0.01, d = -1.15). In follow-up of those results, we explored whether the discrimination index was significantly different from 0 for each group separately. Regardless of shock intensity, the discrimination index was not significantly different from 0 for rats that underwent CS extinction, suggesting that extinction generalized well from CS to GS (MSCS: t(15) = -0.66, p = 0.516, d = -0.17; SSCS: t(15) = 1.658, p = 0.118, d = 0.41). In contrast, the discrimination index was significantly different from 0 for rats that underwent GS extinction (MSGS: t(15) = 7.95, p = < 0.001, d = 1.99; SSGS: t(15) = 3.665, p = 0.002, d = 0.92), indicating that GS extinction did not fully generalize to the CS. One might argue that the calculation of the discrimination index as a relative measure might obscure differences in generalization between the MSGS and SSGS groups, given the higher degree of freezing to the GS in the latter than in the former group. However, an alternative (non-preregistered) measure of discrimination, calculated as the absolute difference in freezing between CS and GS, yielded no difference in discrimination between the MSGS and SSGS groups either (t(60) = -0.695, p = 0.8987, d = -0.25), with discrimination again significantly different from 0 in both groups (MSGS: t(15) = 4.7, p < 0.001, d = 1.17; SSGS: t(15) = 3.39, p = 0.004, d = 0.85).

Further investigation of the significant group by sex interaction revealed differential freezing between sexes in the SSCS group only (t(112) = -3.49, p < 0.001, d = -1.23), with males showing higher freezing than females. Given the high levels of freezing by the male rats of the SSCS group, we investigated if these differences were already present before tone onset. While we observed that contextual freezing levels increased over time (F(4, 224) = 5.39, p < 0.001, ω_p_^2^ = 0.02), overall contextual freezing levels were low (see Figure S3). We observed differences between groups (F(3, 56) = 2.84, p = 0.046, ω ^2^ = 0.02, see Supplementary Table 3), but these differences did not withstand multiple testing correction.

To further evaluate differences in GS extinction retention as a potential source for the differences in CS responding between the SSGS and MSGS group, we compared the difference in freezing between the last two tones of the extinction session and the relevant extinction retention test tones between groups that received the same extinction training (i.e., comparing SSCS to MSCS in their change in CS responding from the end of extinction training to the extinction retention test and SSGS to MSGS in their change in GS responding from the end of extinction training to the extinction retention test). We observed that the difference between the SSGS and MSGS groups in the increase in freezing between both sessions failed to reach significance (F(1, 28) = 3.78, p = 0.062, ω_p_^2^ = 0.08), although it was nominally larger in the SSGS group (M = 12.2, SE = 4.59) than in the MSGS (M = 2.11, SE = 2.43). There was no difference in the increase in freezing from the end of extinction to the extinction retention test when comparing the SSCS and MSCS groups (F(1, 28) = 2.58, p = 0.119, ω_p_^2^ = 0.05).

### Return of fear tests

We presented unsignaled USs, followed by a reinstatement test session 24 hours later (see Figure 5). While we did not observe significant group differences (F(3, 45) = 1.48, p = 0.231, ω_p_^2^ = 0.03), we observed significantly higher freezing to the CS than to the GS (F(1, 45) = 18.1, p < 0.001, ω_p_^2^ = 0.07). No group differences were found in freezing discrimination indices either (F(3, 45) = 1.62, p = 0.199, ω_p_^2^ = 0.04).

**Figure 5.**
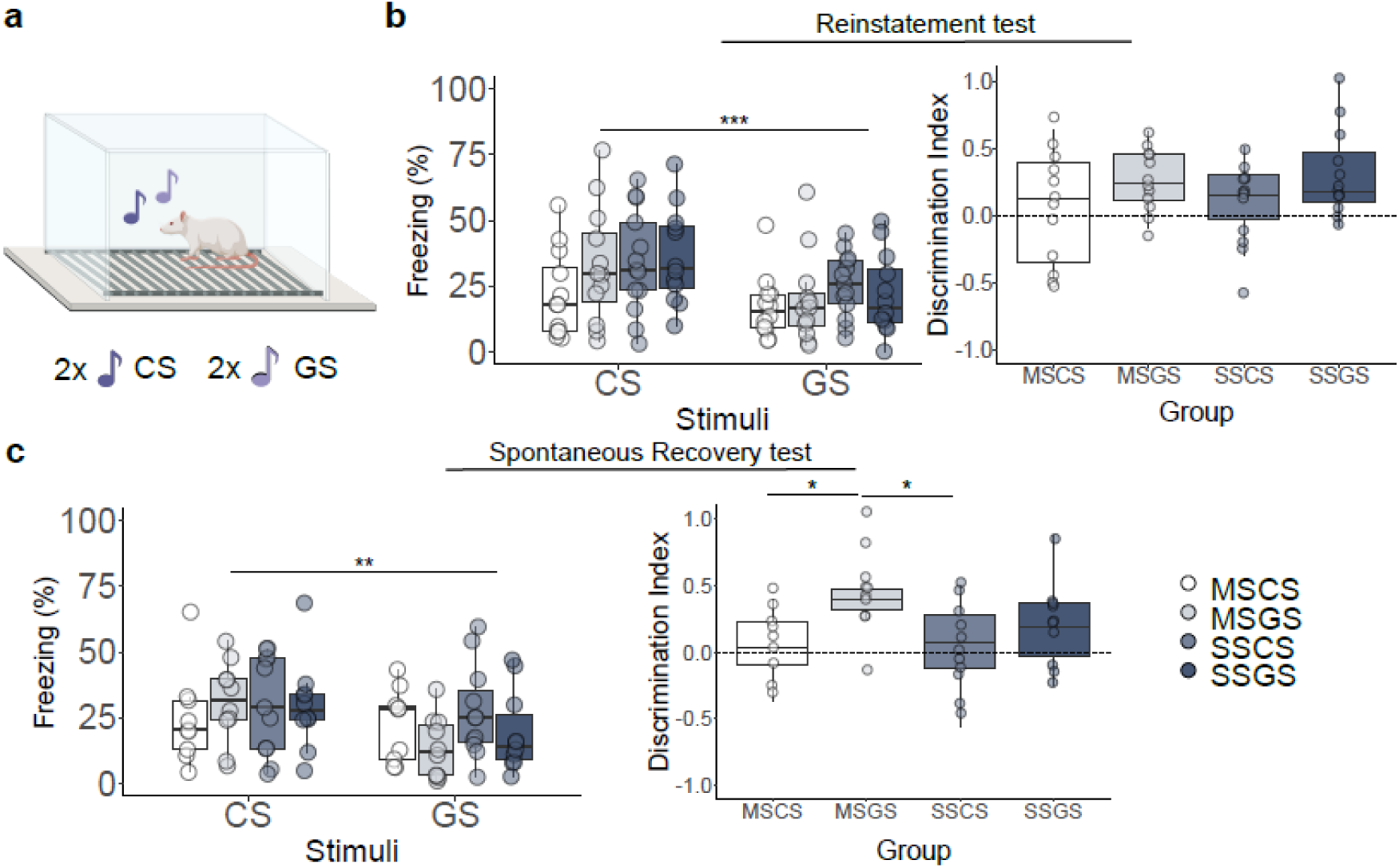
Freezing during reinstatement test and spontaneous recovery test. The freezing boxplots represent the average for each stimulus; results are expressed in % of tone duration. The discrimination index boxplots represent the average freezing discrimination index. Discrimination indices were established by calculating the difference in freezing to the tones (CS minus GS) and dividing by the sum of freezing to both tones. **a.** Graphical representation of the reinstatement test and the spontaneous recovery test. **b.** Results of the reinstatement test session. Significant differences between CS and GS (F(1, 45) = 18.1, p < 0.001) in freezing behavior, but no significant differences in discrimination index between groups were found (F(3, 45) = 1.62, p = 0.199). **c.** Results of the spontaneous recovery test session. Significant differences between CS and GS were found in freezing behavior (F(1, 36) = 10.54, p = 0.003), as well as significant differences in the discrimination index between groups (F(3, 36) = 3.77, p = 0.019).

Seven days later, we tested differences in spontaneous recovery, and found higher freezing to the CS than to the GS again (F(1, 36) = 10.54, p = 0.003, ω ^2^ = 0.06), in the absence of group differences (F(3, 36) = 0.4, p = 0.757, ω_p_^2^ = -0.05) (see Figure 5). We did find group differences in the freezing discrimination index (F(3, 36) = 3.77, p = 0.019, ω ^2^ = 0.17). Specifically, rats in the MSGS group showed a significantly higher discrimination index than those in the MSCS group (t(36) = -2.78, p = 0.041, d = -1.28) and the SSCS group (t(36) = 2.99, p = 0.024, d = 1.31).

## Discussion

We investigated how increased generalization of acquisition affects the generalization of extinction in a Pavlovian fear conditioning task. To achieve differences in generalization of acquisition, in line with previous literature (Laxmi et al., 2003; Xuan et al., 2023), we subjected rats to cued fear conditioning with strong versus moderate intensity shocks as US. While rats that received moderate shocks during acquisition discriminated between the CS and a GS on a subsequent retention test, rats that received strong shocks did not and showed similar levels of freezing to both tones.

In the extinction test session, we confirmed some of our preregistered hypotheses. In the moderate shock groups, we observed the expected asymmetry: in the MSGS group, we observed a high discrimination index, with higher freezing to the CS than to the GS, meaning that extinction did not strongly generalize from GS to CS, while in the MSCS group, the discrimination index was low, in other words, in the MSCS group extinction generalized strongly from the CS to the GS, yielding similar freezing for both tones. In the strong shock groups, we expected that the enhanced generalization of acquisition would further reduce the generalization of extinction, yielding larger discrimination at test between CS and GS particularly in the SSGS group. In support of our hypothesis, we found that animals in the SSGS group showed significantly higher freezing to the CS than those in the MSGS group. We did not observe higher freezing to the GS in the SSCS group than in the MSCS group, which suggests that the enhanced generalization of acquisition from CS to GS as a result of high US intensity went hand-in-hand with reduced generalization of extinction from an extinguished GS back to the CS but not from the extinguished CS to a GS. However, we also noted a tendency for higher responding to the GS in the SSGS group compared to the MSGS group. While not significant after multiple-test correction, this suggests that there was less retention of GS extinction in the SSGS group than in the MSGS group. Hence, it is possible that the SSGS group shows higher responding to the CS than the MSGS group after GS extinction not because of weaker generalization from the GS to the CS but because of the weaker retention of GS extinction leaving less to generalize to the CS. Further research should aim to disentangle these possibilities.

Regarding the return of fear tests, we did not find any group differences in the reinstatement test. We did find that rats showed higher CS freezing than GS freezing, indicative of a selective return of fear to the CS, regardless of initial US intensity or the nature of the stimulus used for extinction training. One week later, we performed a spontaneous recovery test and again observed higher CS-elicited than GS-elicited freezing. Not all our preregistered differences were confirmed here. One reason could be a reduction of power relative to the extinction test session, as some animals were euthanized in between to collect their brain tissue.

It is important to note that the results for the discrimination indices should be interpreted with caution. Rats that underwent GS extinction had mostly received reinforced CS presentations, while the GS was never paired with shock. However, in the groups that underwent CS extinction, the CS became effectively ambiguous, as the CS was paired with shock in acquisition and then not followed by shock during extinction training. Thus, after extinction training, the results of discrimination indices may be biased towards increased discrimination in the MSGS and SSGS groups compared to the MSCS and SSCS groups.

Our data show that the greater generalization that is observed after acquisition with a strong-intensity US is accompanied by stronger retention of CS-elicited fear upon GS extinction. Our results do not allow to discern to what degree this is due to a reduction in generalization of extinction from GS to CS proper, as hypothesized, or to an impaired retention of GS extinction in rats trained with strong-intensity shocks. Future research is needed to dissect this. Of note, the effect did not extend to rats that underwent CS extinction, whose freezing towards the GS at test was the same whether trained with moderate- or high-intensity shocks.

While future studies might seek to replicate and elaborate these results in humans, it would also be worthwhile to investigate the neurobiological mechanisms involved in and constraining the generalization of extinction or the retention of GS extinction in further animal work. Whereas proactively limiting the generalization of fear after a threatening event might be difficult to achieve, finding ways to improve the generalization of extinction might be more feasible. This could prove beneficial then as an adjunct to exposure therapy. As one avenue for further exploration, previous research has shown that vagus nerve stimulation during extinction can increase generalization of extinction in male rats (Noble et al., 2019; Souza et al., 2020).

The use of equal numbers of male and female animals allowed us to explore sex differences. We detected such differences, especially in acquisition and extinction, although less pronounced than expected. We compared freezing between an early extinction block (i.e., first three tones) and a late block (i.e., last three tones), which revealed that male rats showed higher freezing in early and late trials than female rats during those same trials. Such generally reduced freezing in female rats during extinction training has been reported previously in other tasks (Shanazz et al., 2022). While we also expected slower extinction in female rats based on prior literature (Baran et al., 2009; Fenton et al., 2016; Graham et al., 2009; Greiner et al., 2019), we found no supporting evidence for this in our results.

Surprisingly, males in the SSCS group showed high freezing to the CS despite having undergone extinction training with that same CS in the preceding session. This result is puzzling given that their extinction training results look as expected, exhibiting a gradual reduction in freezing over extinction trials. Thus, it seems that males in the SSCS group showed poor extinction retention during the extinction test session. This result could not be attributed to increased contextual fear, nor to technical issues, as rats from all experimental groups were tested simultaneously.

The lack of sex differences during the extinction test was unexpected, as we anticipated lower discrimination indices in female rats, in line with previous reports (Greiner et al., 2019; Keiser et al., 2017). Note that Keiser et al. (2017) assessed generalization of contextual rather than cued fear. While Greiner et al. (2019) found a lack of extinction to a conditioned fear cue and a lack of discrimination between a conditioned fear cue and a conditioned safety cue in females when assessing freezing behavior, females in that study did show differential darting behavior in response to a safety versus a fear cue.

Darting behavior has previously been introduced as a conditioned behavior expressed largely by female rats (Gruene et al., 2015; Mitchell et al., 2022), although the associative properties of darting have been challenged (Trott et al., 2022). In line with previous research (Gruene et al., 2015; Mitchell et al., 2022), we observed that females were more likely to be classified as darters than males (47% compared to 19%). We also found that during acquisition, female darters showed less freezing than female non-darters, but this difference was not significant in their male counterparts nor in any of the other experimental sessions. Two important sidenotes are that female darters darted more than male darters, and that, whereas our female sample had a well-balanced distribution of darters versus non-darters (15 darters vs 17 non-darters), the distribution was very unbalanced for the males (6 darters vs 26 non-darters), which makes these statistical comparisons more unreliable in male rats.

In conclusion, we found higher responding to the CS after GS extinction in strong shock rats compared to moderate shock rats. This could be either due to weaker retention of GS extinction in strong shock rats (while CS extinction is not similarly affected) or due to weaker generalization, as hypothesized. This may be important to consider in extinction-based exposure therapy, as patients frequently exhibit broadly generalized fears, and exposure treatment typically involves cues or situations different from those to which the fear was originally acquired.

### Research data statement

The datasets generated and/or analyzed during the current study are available on the OSF repository: http://doi.org/10.17605/osf.io/9yd83 (Lopez-Moraga et al., 2023).

## Acknowledgements

We thank Yana van der Heyden for assistance with behavioral scoring. Graphical representations of the behavioral setups in the figures were made with Biorender.

## Author contributions

A. L-M.: conceptualization, data curation, formal analysis, funding acquisition, investigation, methodology, project administration, software, validation, visualization, writing – original draft. Z. G.: data curation, investigation; writing – review & editing. L. L.: conceptualization, funding acquisition, methodology, project administration, resources, supervision, writing – review & editing. T. B.: conceptualization, methodology, funding acquisition, project administration, resources, supervision, writing – review & editing.

