## Supplemental figures and tables for "High threat intensity increases conditioned fear generalization and makes extinction of a generalization stimulus less effective"

Supplementary files

Supplementary Figures

**Supplementary Figure 1**

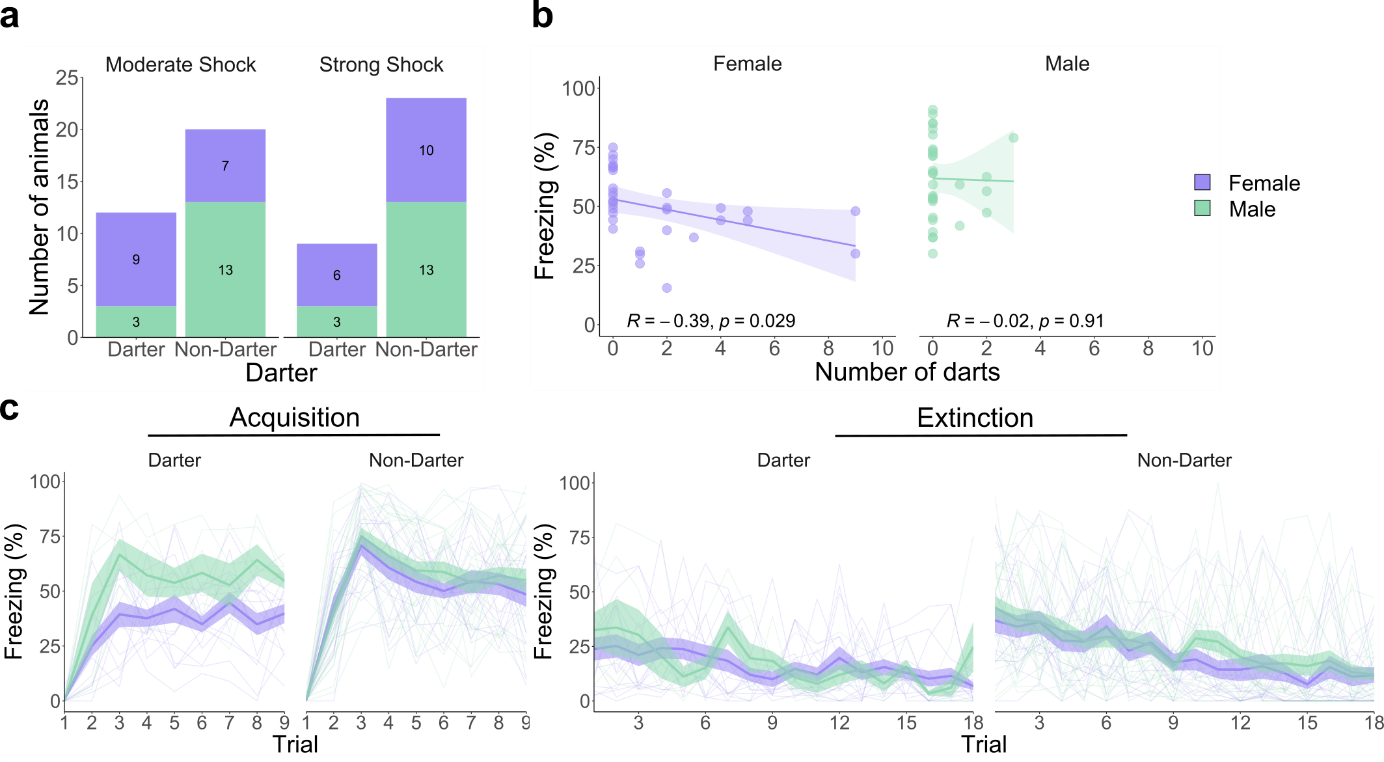

**Figure S1.** Darting analyses during acquisition (a-c) and extinction (d). **a.** Bar plot showing the distribution of darter and non-darter rats across sex and moderate versus strong shock groups. **b.** Correlation plot of freezing and number of darts during CS3 to CS7. Females show a negative correlation between freezing and number of darts (R = -0.39, p = 0.029), while males do not (R = -0.02, p = 0.91). **c.** Female darter rats showed lower freezing than female non-darters (t(30) = -4.8, p < 0.001) during acquisition, while in males there were no differences in freezing between darters and non-darters (t(30) = -0.34, p = 0.73). **d.** Darters and non-darters showed similar levels of freezing during extinction, regardless of sex (females: t(30) = -1.37, p = 0.181; males: t(30) = -1.3, p = 0.202).

**Supplementary Figure 2**

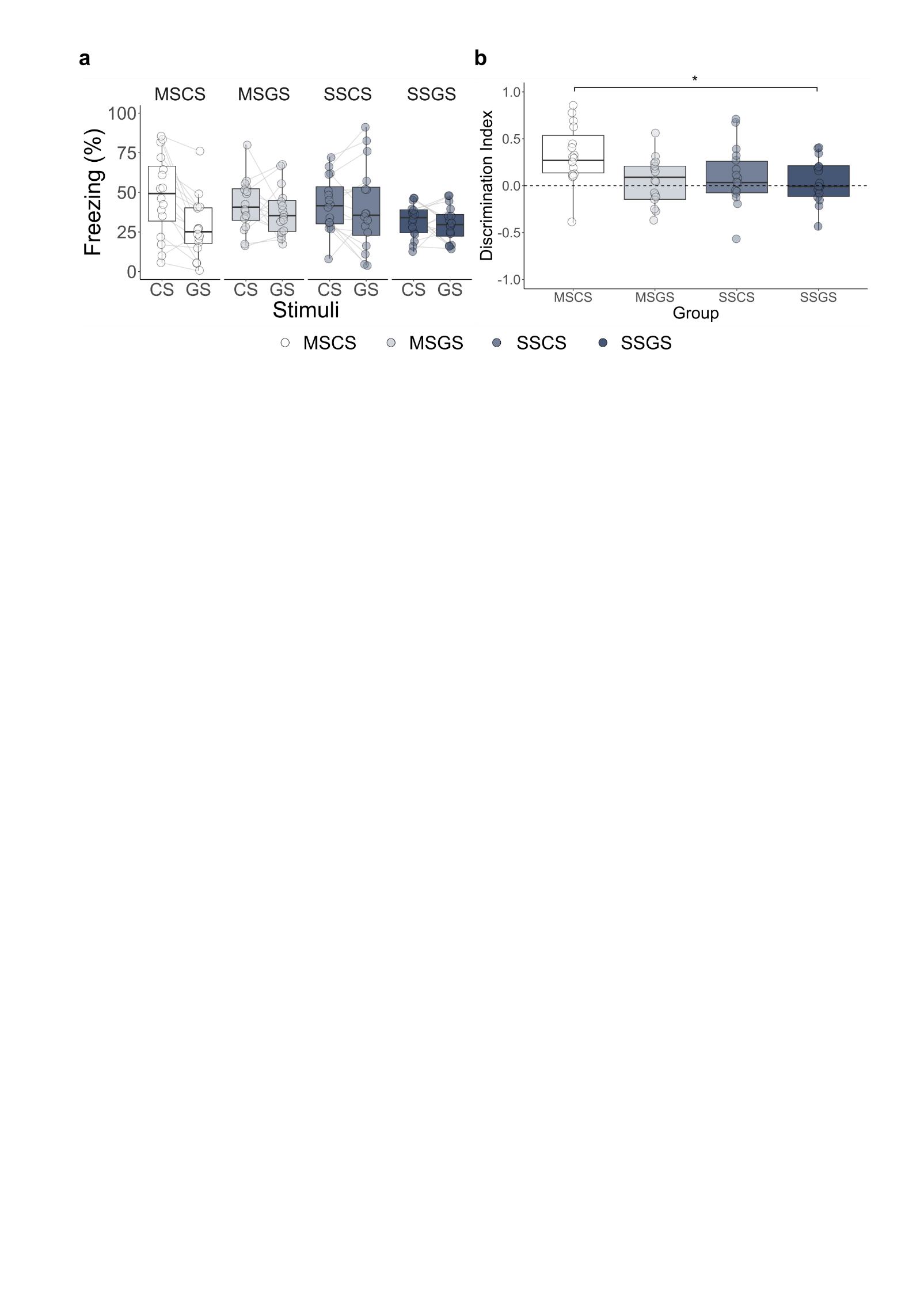

**Figure S2.** Results of the acquisition test session in all groups. Freezing boxplots represent average freezing to each stimulus type, expressed as % of tone duration. Discrimination boxplots represent the average freezing discrimination index per group. Discrimination indices were calculated as the difference in freezing during the presentation of the CS versus the GS, divided by the sum of freezing to both tones. **a.** A significant interaction between group and stimulus was detected (F(3, 60) = 4.21, p = 0.009), but further examination of the interaction did not yield any significant effects (e.g., animals in group SSGS did not show significantly more freezing to the CS than animals in group MSGS). **b.** Discrimination indices were different between the groups (F(3, 60) = 3.17, p = 0.031). In particular, the MSCS group showed a higher discrimination index than the SSGS group (t(60) = 2.73, p = 0.041).

**Supplementary Figure 3**

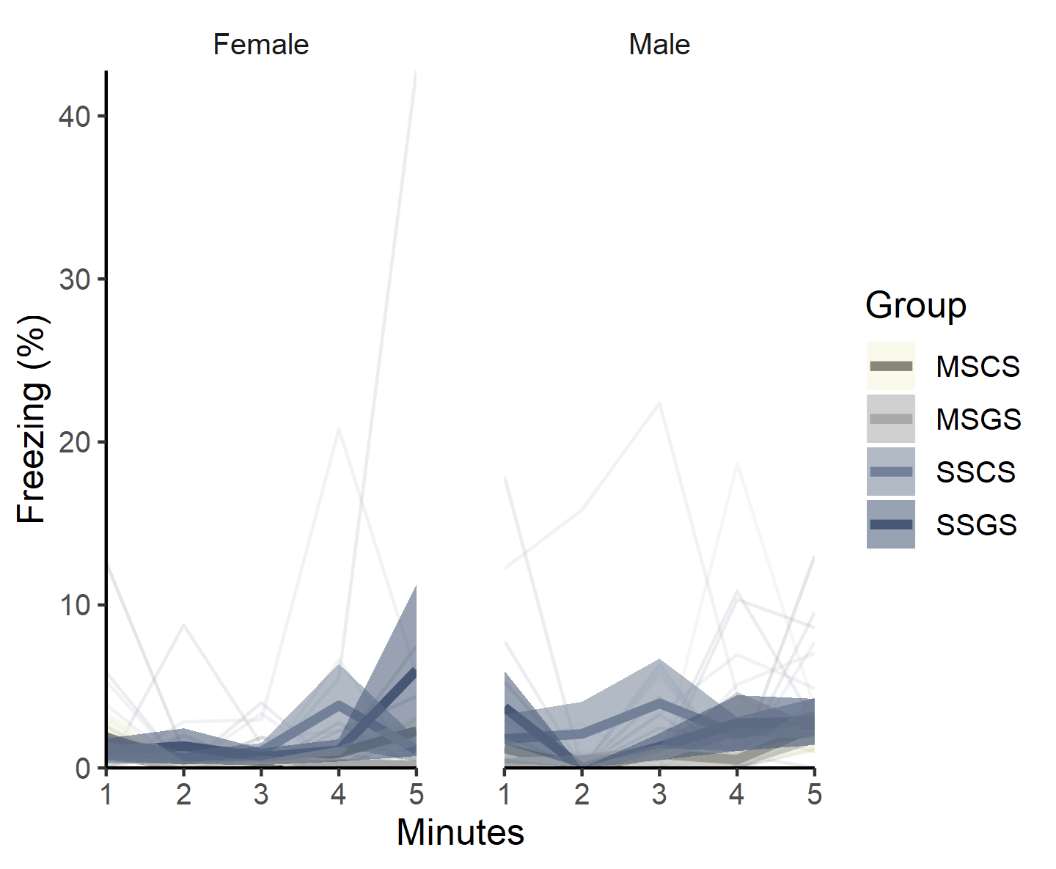

**Figure S3.** Contextual freezing during the first 5 minutes of the extinction test session. Freezing increased over the five minutes before the first tone (F(4, 224) = 5.39, p < 0.001), but was overall low (1.5% ± 0.22). A significant difference in freezing was observed between groups (F(3, 56) = 2.84, p = 0.046), but post hoc analyses yielded no significant effects.

Supplementary Tables

**Supplementary Table 1.** Statistical test results of darting and freezing analyses

| **Phase** | **Measured outcome** | **Statistical test** | **Result** | **Effect size** |
| --- | --- | --- | --- | --- |
| Acquisition | Darting and Sex association | Chi square test | χ^2^(1)= 4.54, **p = 0.033** | φ = 0.3 [0, 1] |
| Acquisition | Darting and Group shock association | Chi square test | χ^2^(1)= 0.28, p = 0.594 | φ = 0.1 [0.09, 1] |
| Acquisition | Darting and Group association | Chi square test | χ^2^(3)= 5.88, p = 0.117 | Cramer’s V = 0.21 [0, 1] |
| Acquisition | Freezing - Females | Two-sided t-test | Darter vs Non-darter: t(30) = -4.8, **p < 0.001** | d = -1.7 [-2.44, -1.14] |
| Acquisition | Freezing - Males | Two-sided t-test | Darter vs Non-darter: t(30) = -0.34, p = 0.73 | d = -0.17 [-1.07, 0.71] |
| Acquisition test | Freezing - Females | Two-sided t-test | Darter vs Non-darter: t(30) = 0.18, p = 0.857 | d = 0.06 [-0.63, 0.76] |
| Acquisition test | Freezing - Males | Two-sided t-test | Darter vs Non-darter: t(30) = -0.38, p = 0.71 | d = -0.17 [-1.06, 0.72] |
| Extinction training | Freezing - Females | Two-sided t-test | Darter vs Non-darter: t(30) = -1.37, p = 0.181 | d = -0.49 [-1.21, 0.26] |
| Extinction training | Freezing - Males | Two-sided t-test | Darter vs Non-darter: t(30) = -1.3, p = 0.202 | d = -0.64 [-1.85, 0.3] |
| Extinction test | Freezing - Females | Two-sided t-test | Darter vs Non-darter: t(30) = -0.24, p = 0.808 | d = -0.09 [-0.78, 0.61] |
| Extinction test | Freezing - Males | Two-sided t-test | Darter vs Non-darter: t(30) = -0.27, p = 0.791 | d = -0.12 [-1.01, 0.77] |
| Reinstatement test | Freezing - Females | Two-sided t-test | Darter vs Non-darter: t(23) = -0.26, p = 0.798 | d = -0.1 [-0.89, 0.68] |
| Reinstatement test | Freezing - Males | Two-sided t-test | Darter vs Non-darter: t(22) = -0.47, p = 0.646 | d = -0.23 [-1.22, 0.76] |
| Spontaneous recovery test | Freezing - Females | Two-sided t-test | Darter vs Non-darter: t(18) = -1.04, p = 0.312 | d = -0.47, [-1.36, 0.43] |
| Spontaneous recovery test | Freezing - Males | Two-sided t-test | Darter vs Non-darter: t(18) = -0.45, p = 0.658 | d = -0.23 [-1.24, 0.79] |

**Supplementary Table 2.** Additional statistical test results of freezing to the CS and GS and discrimination during the acquisition test

| **Measured outcome** | **Statistical test** | **Result** | **Effect size** |
| --- | --- | --- | --- |
| Freezing | Mixed ANOVA | Group: F(3, 60) = 1.18, p = 0.324  Stimuli: F(1, 60) = 10.89, **p = 0.002**  Group*Stimuli: F(3, 60) = 4.21, **p = 0.009** | ω_p_^2^ < 0.01 [0, 1]  ω_p_^2^ =0.03 [0, 1]  ω_p_^2^ = 0.03 [0,1] |
| Freezing | Simple effects by stimuli with Tukey correction - CS | MSCS – MSGS: t(120) = 0.909, p = 0.8  MSCS – SSCS: t(120) = 0.898, p = 0.806  MSCS – SSGS: t(120) = 2.441, p = 0.075  MSGS – SSCS: t(120) = -0.011, p = 1  MSGS – SSGS: t(120) = 1.531, p = 0.42  SSCS – SSGS: t(120) = 1.542, p = 0.416 | d = 0.32 [-0.38, 1.02]  d = 0.32 [-0.38, 1.01]  d = 0.86 [0.15, 1.57]  d < -0.01 [-0.7, 0.69]  d = 0.54 [-0.16, 1.24]  d = 0.54 [-0.16, 1.25] |
| Freezing | Simple effects by stimuli with Tukey correction - GS | MSCS – MSGS: t(120) = -1.575, p = 0.396  MSCS – SSCS: t(120) = -1.896, p = 0.235  MSCS – SSGS: t(120) = 0.410, p = 0.977  MSGS – SSCS: t(120) = -0.320, p = 0.989  MSGS – SSGS: t(120) = 1.165, p = 0.65  SSCS – SSGS: t(120) = 1.485, p = 0.449 | d = -0.55 [-1.26, 0.15]  d = -0.67 [-1.38, 0.03]  d = -0.14 [-0.84, 0.55]  d = -0.11 [-0.81, 0.59]  d = 0.41 [-0.29, 1.11]  d = 0.52 [-0.18, 1.23] |
| Discrimination index | One-way ANOVA | Group: F(3, 60) = 3.17, **p = 0.031** | ω_p_^2^ =0.09 [0, 1] |
| Discrimination index | Pairwise comparisons with Tukey correction | MSCS – MSGS: t(60) = 2.58, p = 0.06  MSCS – SSCS: t(60) = 2.01, p = 0.169  MSCS – SSGS: t(60) = 2.73, **p = 0.041**  MSGS – SSCS: t(60) = -0.57, p = 0.94  MSGS – SSGS: t(60) = 0.15, p = 0.999  SSCS – SSGS: t(60) = 0.72, p = 0.889 | d = 0.91 [0.19, 2.64]  d = 0.71 [-0.01, 1.43]  d = 0.96 [0.24, 1.69]  d = -0.2 [-0.91, 0.51]  d = 0.05 [-0.66, 0.76]  d = 0.25 [-0.45, 0.96] |

**Supplementary Table 3.** Statistical results of contextual freezing during the extinction test

| **Measured outcome** | **Statistical test** | **Result** | **Effect size** |
| --- | --- | --- | --- |
| Contextual freezing | Non-parametric mixed ANOVA | Sex: F(1, 56) = 0.49, p = 0.484  Group: F(3, 56) = 2.84, **p = 0.046**  Minute: F(4, 224) = 5.39, **p < 0.001**  Sex*Group: F(3, 56) = 0.29, p = 0.832  Sex*Minute: F(4, 224) = 0.97, p = 0.424  Group*Minute: F(12, 224) = 0.65, p = 0.797  Sex*Group*Minute: F(12, 224) = 0.97, p = 0.48 | ω_p_^2^ < 0.01 [0, 1]  ω_p_^2^ = 0.02 [0, 1]  ω_p_^2^ = 0.02 [0,1]  ω_p_^2^ = -0.04 [0,1]  ω_p_^2^ = -0.01 [0,1]  ω_p_^2^ = -0.01 [0,1]  ω_p_^2^ < -0.01 [0,1] |
| Contextual freezing | Pairwise comparisons with Tukey correction | MSCS – MSGS: t(56) = 1.18, p = 0.642  MSCS – SSCS: t(56) = -1.23, p = 0.613  MSCS – SSGS: t(56) = -1.37, p = 0.525  MSGS – SSCS: t(56) = -2.41, p = 0.087  MSGS – SSGS: t(56) = -2.55, p = 0.064  SSCS – SSGS: t(56) = -0.14, p = 0.999 | r = 0.08 [-0.1, 0.25]  r = -0.09 [-0.26, 0.09]  r = -0.03 [-0.21, 0.14]  r = -0.15 [-0.32, 0.02]  r = -0.1 [-0.28, 0.08]  r = 0.04 [-0.13, 0.22] |
